# Long non-coding RNAs defining major subtypes of B cell precursor acute lymphoblastic leukemia

**DOI:** 10.1101/365429

**Authors:** Alva Rani James, Michael P Schroeder, Martin Neumann, Lorenz Bastian, Cornelia Eckert, Nicola Gökbuget, Jutta Ortiz Tanchez, Cornelia Schlee, Konstandina Isaakidis, Stefan Schwartz, Thomas Burmeister, Arend von Stackelberg, Michael A Rieger, Stefanie Göllner, Martin Horstman, Martin Schrappe, Renate Kirschner-Schwabe, Monika Brüggemann, Carsten Müller-Tidow, Hubert Serve, Altuna Akalin, Claudia D Baldus

## Abstract

Recent studies implicated that long non-coding RNAs (lncRNAs) may play a role in the progression and development of acute lymphoblastic leukemia, however, this role is not yet clear. In order to unravel the role of lncRNAs associated with B-cell precursor Acute Lymphoblastic Leukemia (BCP-ALL) subtypes, we performed transcriptome sequencing and DNA methylation array across 82 BCP-ALL samples from three molecular subtypes (DUX4, Ph-like, and Near Haploid or High Hyperdiploidy). Unsupervised clustering of BCP-ALL samples on the basis of their lncRNAs on transcriptome and DNA methylation profiles revealed robust clusters separating three molecular subtypes. Using extensive computational analysis, we developed a comprehensive catalog of 1235 aberrantly dysregulated BCP-ALL subtype-specific lncRNAs with altered expression and methylation patterns from three subtypes of BCP-ALL. By analyzing the co-expression of subtype-specific lncRNAs and protein-coding genes, we inferred key molecular processes in BCP-ALL subtypes. A strong correlation was identified between the DUX4 specific lncRNAs and activation of TGF-β and Hippo signaling pathways. Similarly, Ph-like specific lncRNAs were correlated with genes involved in activation of PI3K-AKT, mTOR, and JAK-STAT signaling pathways. Interestingly, the relapse-specific differentially expressed lncRNAs correlated with the activation of metabolic and signaling pathways. Finally, we showed a set of epigenetically altered lncRNAs facilitating the expression of tumor genes located at their *cis* location. Overall, our study provides a comprehensive set of novel subtype and relapse-specific lncRNAs in BCP-ALL. Our findings suggest a wide range of molecular pathways are associated with lncRNAs in BCP-ALL subtypes and provide a foundation for functional investigations that could lead to new therapeutic approaches.

**Author Summary:** Acute lymphoblastic leukemia is a heterogeneous blood cancer, with multiple molecular subtypes, and with high relapse rate. We are far from the complete understanding of the rationale behind these subtypes and high relapse rate. Long non-coding (lncRNAs) has emerged as a novel class of RNA due to its diverse mechanism in cancer development and progression. LncRNAs does not code for proteins and represent around 70% of human transcripts. Recently, there are a number of studies used lncRNAs expression profile in the classification of various cancers subtypes and displayed their correlation with genomic, epigenetic, pathological and clinical features in diverse cancers. Therefore, lncRNAs can account for heterogeneity and has independent prognostic value in various cancer subtypes. However, lncRNAs defining the molecular subtypes of BCP-ALL are not portrayed yet. Here, we describe a set of relapse and subtype-specific lncRNAs from three major BCP-ALL subtypes and define their potential functions and epigenetic regulation. Our data uncover the diverse mechanism of action of lncRNAs in BCP-ALL subtypes defining how lncRNAs are involved in the pathogenesis of disease and the relevance in the stratification of BCP-ALL subtypes.

## INTRODUCTION

B-cell Precursor Acute Lymphoblastic Leukemia (BCP-ALL) is the most prevalent disease in children and affects also adults. Despite improvements in treatment regimens such as chemotherapy and allogeneic hematopoietic stem cell transplantation, the prognosis remains poor for patients in high-risk groups and at relapse (1). Various risk subtypes have been established based on the cytogenetic analysis and molecular genetics studies. These subtypes are classified based on the presence of high hyperdiploidy (51-65 chromosomes)(2), hypodiploidy (less than 44 chromosomes)(3) and fusion genes (for example BCR-ABL, ETV6-RUNX, MLL, etc)(4). About 70-80% of both adults and pediatric cases of BCP-ALL constitute these subtypes, although the frequency may differ (5).

Recent efforts taking advantage of whole transcriptome sequencing (RNA-Seq) have refined this classification by identifying novel BCP-ALL subtypes. RNA-Seq analysis identified cytogenetically non-detectable recurrent rearrangements and gene fusions, which allowed characterization of additional subtypes based on distinct gene expression profiles (6). For example, the DUX4 (7) subtype is defined mainly by the IGH-DUX4 or ERG-DUX4 gene fusions; the Ph-like (8) subtype is a high-risk subtype with a gene expression profile similar to Ph-positive ALL; however, lacking BCR-ABL1 fusion gene; and the Near Haploid/High Hyperdiploid (NH-HeH) (51–67 chromosomes) subtype (9,10) is a common subtype, comprising 30% of all pediatric BCP-ALL. These subtypes are clinically relevant with distinct gene expression profile and have been widely studied in the recent past. Nevertheless, we are far from complete understanding of BCP-ALL subtypes and their heterogeneity and its associated molecular mechanisms, which pose a major challenge for improving diagnosis and therapy. Recent studies have suggested that long non-coding RNAs (lncRNAs) and small non-coding RNAs (e.g. microRNAs) might play a key role in development and progression of leukemia (11) and thus constitute as new biomarkers and potential targets for novel therapies (12).

LncRNAs are arbitrarily defined as transcripts longer than 200 base pairs and lacking an extended protein-coding open reading frame (ORF). It has become apparent that lncRNAs are frequently spliced and polyadenylated and are mainly transcribed by RNA polymerase II (13). LncRNAs expression has been reported as highly tissue-specific even though the expression abundance is generally lower compared to protein-coding genes (14). The expression specificity has been extended to a wide variety of physiological and pathological mechanisms like cancer development and Pluripotency (15). LncRNAs can act either proximally (in the cis region) or distally (in the trans region) for the transcriptional regulation of protein-coding genes (16). Like proteins, various lncRNAs are attributed to oncogenic or tumor-suppressive (17) activities exerting various cellular functions (18). In addition, lncRNAs regulate gene expression at the epigenetic (19) and post-transcription (20) levels. Genome-wide association studies in cancer have disclosed that 80% of cancer-associated single-nucleotide polymorphisms (SNPs) are in non-coding regions (21), including lncRNAs, suggesting that a significant portion of the genetic etiology of cancer can be related to lncRNAs (22). Moreover, lncRNAs are reported to be useful for disease prognosis, exemplified by the lncRNA HOTAIR (HOX transcript antisense RNA), which is up-regulated in acute myeloid leukemia (AML) patients (23). So far, the majority of studies explored the role of single lncRNAs in leukemia including AML (24), chronic lymphocytic leukemia (CLL) (25) and pediatric ALL (26). Yet a comprehensive genomic and epigenetic delineation of lncRNAs deregulations in BCP-ALL subtypes, and their molecular and functional insights are lacking.

In the present study, we explored lncRNA landscapes in DUX4, Ph-like, and NH-HeH BCP-ALL subtypes and extracted novel biological and functional insights of BCP-ALL subtype-specific lncRNAs and their epigenetic activity. On the basis of RNA-seq transcriptional and DNA methylation survey of lncRNAs, we have determined 1235 subtype-specific, relapse-specific markers and epigenetically altered lncRNAs and demonstrated their relevance in BCP-ALL subtype classification. From our in-depth analyses, we have inferred the potential functions of subtype-specific lncRNAs. Overall, this work provides a most comprehensive and integrative resource which highlights the impact of lncRNAs on relevant pathways that are dysregulated in the molecular subgroups of BCP-ALL and may provide new approaches for prognosis and treatment.

## RESULTS

### Unique lncRNAs expression profiles characterize BCP-ALL subtypes

To identify BCP-ALL subtype-specific lncRNAs, we analysed transcriptome profiles from paired initial diagnosis (ID) and relapse (REL) samples of 26 pediatric and 22 adult BCP-ALL patients lacking known chromosomal translocations like BCR-ABL. Based on DNA mutations and chromosomal translocations combined with RNA expression and DNA methylation profiles the samples were classified into different molecular subtypes (Table S1), namely DUX4 (n = 23), Ph-like (n = 21), Near Haploid or High Hyperdiploid (NH-HeH) (n = 16), and low-hypodiploid (LH) (n = 6) and others (n = 18).

When the distribution of lncRNAs gene expression levels across all BCP-ALL samples was compared with that of protein-coding genes, the former generally showed lower expression levels than the latter (27) (Fig S1A, Table S1). The principal component analysis (PCA) on the expression (FPKM value) of 13,860 GENCODE lncRNAs revealed three major BCP-ALL subtypes, DUX4, Ph-like and NH-HeH with a distinct separation (Fig 1A). This observation is in concordance to the predefined molecular classification. In particular, samples of the DUX4 subtype showed robust separation compared to the remaining samples implying a subtype-specific lncRNAs signatures.

**Fig 1:**
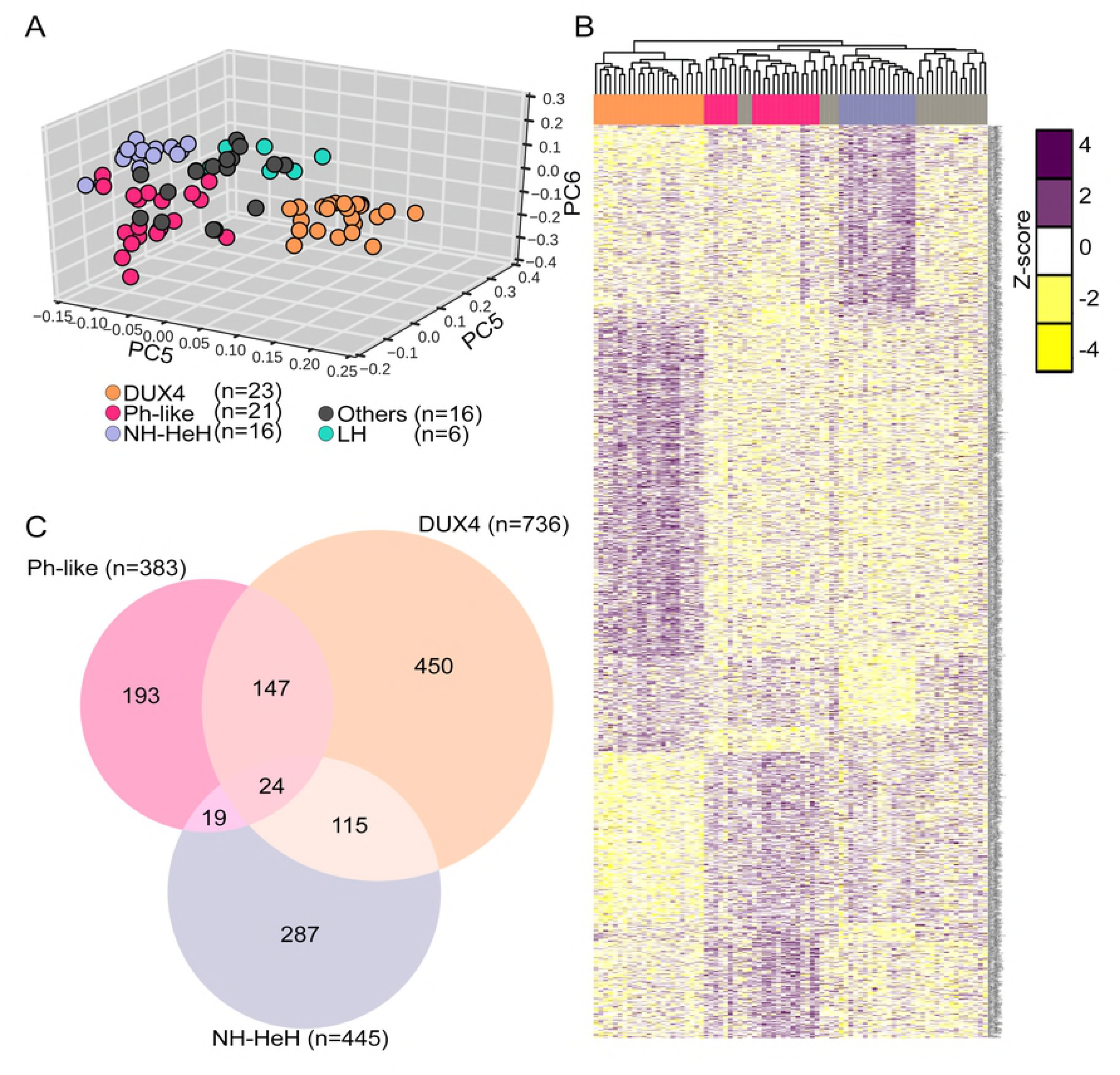
BCP-ALL subtype-specific lncRNA expression signatures. (A) PCA plot constructed from expression FPKM values of lncRNAs from 82 BCP-ALL samples obtained from RNA-seq. Each point represents a BCP-ALL sample. DUX4, Ph-like, NH-HeH, LH subtype and others are represented by orange, rose, blue, green and gray respectively. (B) Heatmap illustrates hireachial clustering DE subtype-specific lncRNAs (absolute Fold change >= +-1.5, *P*-value <= 0.01) signature based on z-score transformed LIMMA normalized expression values on 930 subtype-specific lncRNAs from DUX4 (n = 450), Ph-like (n = 193), and NH-HeH (n = 287) subtypes. (C) The venn diagram illustrates the overlap between subtype-specific lncRNAs from three subtypes, showing 24 lncRNAs are to be common for all three subtypes.

To unveil differentially expressed (DE) lncRNAs across these three major molecular subtypes, we performed DE analysis between subtypes. We obtained 1235 significant DE subtype-specific lncRNAs (*P*-value <= 0.01 and absolute Fold change >= +-1.5) defining signatures of three subtypes (Fig 1B, Fig S2A-C, Table S1). Of these, 24 lncRNAs were commonly detected in all 3 BCP-ALL subtypes (Fig 1C), about 523 (Hypergeometric *P*-value = 9.2E-29) subtype-specific lncRNAs overlapped with deregulated lncRNAs from 12 other cancer types (Fig S2D, Table S1) (28). The remaining 46% (n = 713) of BCP-ALL subtype-specific DE lncRNAs were novel and specific to our subtypes. Out of the overlapped DE lncRNAs (n = 523), 23 (Table S1) were cross-validated in independent cohorts from lnc2cancer (29) database and found to be enriched for oncogenic class of lncRNAs. For example, oncogenic lncRNAs *PVT1* (30) and *GAS5* (31) are differentially up-regulated in the DUX4 subgroup, and *CRNDE* (32) is DE in Ph-like subgroup. Together, subtype-specific lncRNAs signatures assigned molecular subgroups of BCP-ALL.

### Identification and inferred functions of lncRNAs associated with molecular subtypes of BCP-ALL

As lncRNAs can function by regulating protein-coding genes in *cis* and/or *trans* (33–36) regions, we performed functional enrichment analyses using guilt-by-association approach based on the correlation between neighbouring (*cis*) and distally (*trans*) located protein-coding (PC) genes (within ± 100 kb cis and >± 100 kb window for trans) of the subtype-specific lncRNAs (see materials and methods). Expression of both *cis* and *trans* PC genes showed a higher tendency towards positive correlation with the expression of the corresponding lncRNAs (Table 1).

**Table 1:** Number of BCP-ALL subtype-specific lncRNAs co-expressed with its *cis* and *trans* PC genes.

| Subtypes | *Cis* PC genes<br>(n = 929) | *Cis* co-expressed<br>DE lncRNAs<br>(n = 62) | *Trans* PC<br>genes<br>(n = 753) | *Trans* co-expressed<br>DE lncRNAs<br>(n = 552) |
| --- | --- | --- | --- | --- |
| Ph-like | 260 | 170 (383) | 261 | 173 (383) |
| DUX4 | 669 | 451 (736) | 492 | 379 (736) |

The table represents the number of DE lncRNAs showed *cis* ( $\leq 100$ Kb proximity) and *trans* ( $\geq 100$ Kb) protein coding genes and the number of DE lncRNAs co-expressed with them. The numbers shown within the bracket is the total number of DE lncRNAs corresponding to the respective subtypes.

Significantly co-expressed (Pearson correlation coefficient => 0.55, 2-tailed *P*-value <= 0.05) *cis* and *trans* protein-coding genes associated with DUX4 (n = 58 in *cis* and n = 127 in *trans*) and Ph-like (n = 24 in *cis* and n = 20 in *trans*) specific DE lncRNAs demonstrated activation of key signalling pathways involved in proliferation, apoptosis, and differentiation in leukemia (Table S2). For example, in the *cis* based analysis, we identified a strong correlation between DUX4 specific lncRNAs and genes involved in the TGF-beta, Hippo, and P53 signalling pathways (Fig 2A, Table S2). Whereas, the Ph-like specific lncRNAs were correlated with genes involved in JAK-STAT, mTOR, and PIK3-AKT signalling pathways (Fig 2B, Table S2). The *trans* based analysis revealed same vital signalling pathways in DUX4 subtype (Fig S3A-B, Table S2), whereas in Ph-like subtype we identified additional signalling pathways, including, P53 and mitogen-activated protein kinase (MAPK) pathways (Fig S3C, Table S2). The strongly co-expressed *cis* PC genes with DE lncRNAs (n = 32) includes oncogenes including, *IL2RA* (37), *TGFB2* (38), and *CDK6* (39) activated in signalling pathways from DUX4 and Ph-like subgroups (Fig S4A-D, Table 2-3).

**Fig 2:**
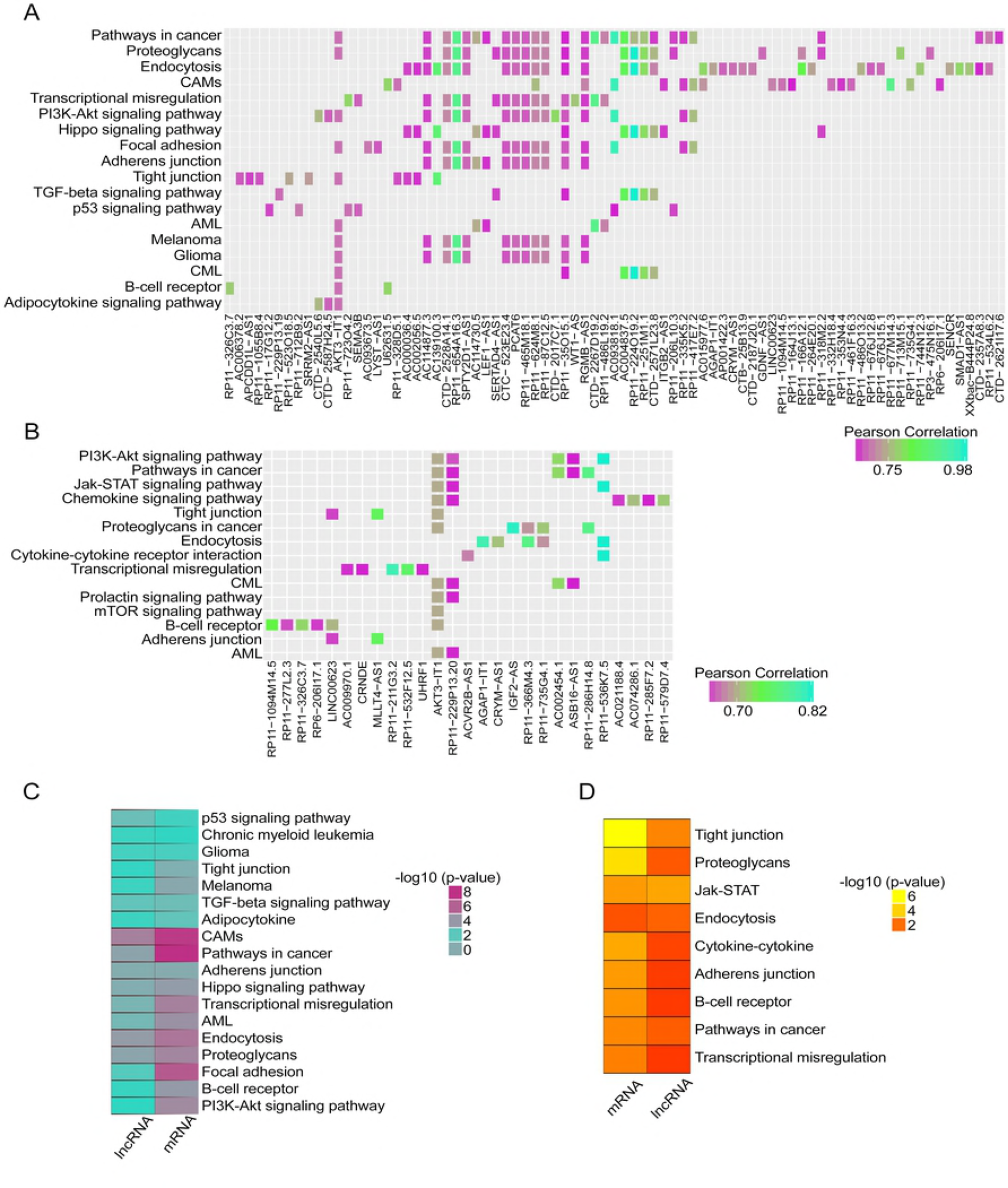
The molecular pathways of lncRNAs involved in the DUX4 and Ph-like BCP-ALL subgroups. (A) The plot depicts the molecular pathway analysis from the functional enrichment analysis for nearby (&3x= 100 kb proximity) cis protein-coding genes correlated (Pearson correlation coefficient >= 0.55 and 2-tailed *P*-value <= 0.05) with DE lncRNAs in the DUX4 subtype. (B) The plot depicts the molecular pathway analysis from the functional enrichment analysis for nearby (<= 100 kb proximity) cis protein-coding genes correlated (Pearson correlation coefficient >= 0.55 and 2-tailed *P*-value <= 0.05) with DE lncRNAs in the Ph-like subtype. (C) The heatmap depicts the concordance between the protein-coding and lncRNAs based predictions for DUX4 subtypes. (D) The heatmap depicts the overlapping pathways from both lncRNAs and protein-coding in the Ph-like subtype. The KEGG pathways or biological functions presented in the heatmaps and barplots show with *P*-value <= 0.05 and > 2 genes involved in each pathways. The hypergeometric p-values are obtained from GeneSCF for the pathways. CAMs : Cell adhesion molecules, CML : Chronic myeloid leukemia, AML : Acute myeloid leukemia.

**Table 2:** Subtype-specific lncRNAs and oncogenes.

| Subtype-specific lncRNAs | Pearson correlation coefficient | P-value | Oncogene |
| --- | --- | --- | --- |
| RP11-347C18.3 | 0.56 | 3.25E-008 | CDK6 |
| RP11-461F16.3 | 0.62 | 5.21E-010 |  |
| RP11-96H19.1 | 0.62 | 3.89E-010 |  |
| RP11-228B15.4 | 0.64 | 7.68E-011 |  |
| MME-AS1 | 0.56 | 3.68E-008 |  |
| CTB-39G8.3 | 0.57 | 1.78E-008 |  |
| AC002454.1 | 0.72 | 2.21E-014 |  |
| RP11-582J16.4 | 0.55 | 8.08E-008 |  |
| AC009970.1 | 0.64 | 6.23E-011 |  |
| RP11-229P13.20 | 0.66 | 1.44E-011 |  |
| LINC00114 | 0.57 | 3.06E-008 |  |
| CTB-118N6.3 | 0.61 | 9.70E-010 |  |
| SOCS2-AS1 | 0.62 | 4.94E-010 |  |
| CTD-2561B21.10 | 0.61 | 9.91E-010 |  |
| RP11-413E1.4 | 0.56 | 4.36E-008 |  |
| KB-1460A1.1 | 0.55 | 7.77E-008 |  |
| AC012309.5 | 0.59 | 4.10E-009 |  |
| RP11-37B2.1 | 0.59 | 4.76E-009 |  |
| ASB16-AS1 | 0.65 | 3.86E-011 |  |
| LINC00426 | 0.62 | 6.32E-010 |  |
| LINC01071 | 0.57 | 2.46E-008 |  |
| RP11-536K7.5 | 0.74 | 5.11E-15 | IL2RA |
| RP11-224O19.2 | 0.98 | 1.08E-061 | TGFB2 |
| AC004837.5 | 0.83 | 6.11E-023 |  |
| RP11-251M1.1 | 0.79 | 7.39E-019 |  |
| CTD-2571L23.8 | 0.75 | 2.94E-016 |  |
| RP11-35O15.1 | 0.65 | 3.36E-011 |  |
| AC139100.3 | 0.58 | 1.00E-008 |  |
| RP11-158M2.3 | 0.58 | 1.50E-008 |  |
| RP11-672A2.5 | 0.56 | 4.68E-008 |  |
| CTD-2357A8.3 | 0.55 | 7.46E-008 |  |
| RP11-677M14.3 | 0.55 | 6.68E-008 |  |

Positively correlating novel *cis* subtype-specific lncRNAs with oncogenes, *CDK6*, *TGFB2*, and *IL2RA* from Ph-like and DUX4 subtypes.

**Table 3:** Subtype-specific novel DE lncRNAs co-expressed with oncogenes, which are associated with vital molecular pathways.

| Subtype-specific lncRNAs | Cis PC | Pearson correlation coefficient | Associated pathways |
| --- | --- | --- | --- |
| RP11-224O19.2<br>AC004837.5<br>RP11-251M1.1<br>CTD-2571L23.8 | TGFB2 | 0.98<br>0.83<br>0.79<br>0.75 | Hippo<br>TGF-β<br>Endocytosis |
| AC093818.1<br>AC078883.3 | ITGA6 | 0.95<br>0.68 | PI3K-Akt |
| U62631.5 | CD22 | 0.78 | Cell adhesion molecules (CAMs)<br>B cell receptor signaling pathway |
| CTD-2267D19.2<br>RP11-486L19.2 | RARA | 0.89<br>0.70 | Pathways in cancer<br>Transcriptional mis-regulation in<br>cancer pathways |

The table represents the novel subtype specific DE lncRNAs co-expressed with its *cis* genes such as *TGFB2*, *ITGA6*, *CD22*, and *RARA* genes, which were enriched in vital molecular pathways in BCP-ALL.

However, there were no significant pathways identified within NH-HeH subtype. The subtype-specific *cis* and *trans* acting lncRNAs which are up-regulated and correlated with genes involved in signalling pathways from DUX4 and Ph-like subtypes were hinting their gene expression regulatory activity.

We next related the functions of DUX4 and Ph-like specific DE lncRNAs obtained from *cis* based analysis to those functions identified with DE PC genes. We observed an overlap of 100% (n = 18, Table S2) of pathways from the DUX4 subtype between lncRNAs based and PC based functional enrichment analysis (Fig 2C). Whereas, in Ph-like subtype, we identified 60% (9 out of 15) same pathways between DE PC based and DE lncRNAs based functional enrichment analysis (Table S2 and Fig 2D). However, we identified Ph-like specific lncRNAs to be more strongly correlated with genes involved in key signalling pathways than Ph-like specific protein-coding genes. For example, we identified mTOR and PI3K-AKT exclusively in the Ph-like specific lncRNAs based analysis. Together, our analyses highlight important functions of BCP-ALL subtype-specific lncRNAs whose expression correlates tightly with that of cancer-related oncogenes.

### Relapse-specific lncRNAs driving BCP-ALL progression

To gain insights into the possible role of lncRNAs driving BCP-ALL progression, we investigated dysregulation of lncRNAs at relapse. For each molecular BCP-ALL subtype, we performed a differential expression analysis of lncRNAs between ID and REL samples (Fig 3). 947 lncRNAs (Table S3) emerged as significantly DE (absolute Fold change >= +-1.5; *P*-value <= 0.01) between ID and REL from the three subtypes. Around 20% (n = 186) of those DE lncRNAs were up-regulated and 80% were down-regulated at relapse. The hierarchical clustering on relapse-specific lncRNAs within each subtype (DUX4, Ph-like, NH-HeH) identified clear separation between ID and REL (Fig 3A-C). We observed 19% (183) relapse-specific lncRNAs identified here overlapped with subtype-specific lncRNAs (Fig 3E). The putative molecular functions of relapse-specific lncRNAs were identified using the previously mentioned guilt-by-association approach. Relapse-specific lncRNAs within Ph-like and NH-HeH subtypes did not show any significant correlation with activation of pathways. In contrast, in the DUX4 subtype, we identified 56% (n = 321) relapse-specific lncRNAs correlated with *cis* PC genes (Table S3). These strongly correlated relapse-specific lncRNAs showed activation of PC genes involved in vital signalling pathways and metabolic pathways, including NF-kappa B-signalling pathway, cell adhesions molecule (CAMS) and metabolic pathways (number of genes involved >= 3 and *P*-value <= 0.05) (Fig 3D, Table S3). These results indicate that relapse-specific markers from DUX4 subtype may be functionally engaged in metabolic and signalling pathways.

**Fig 3:**
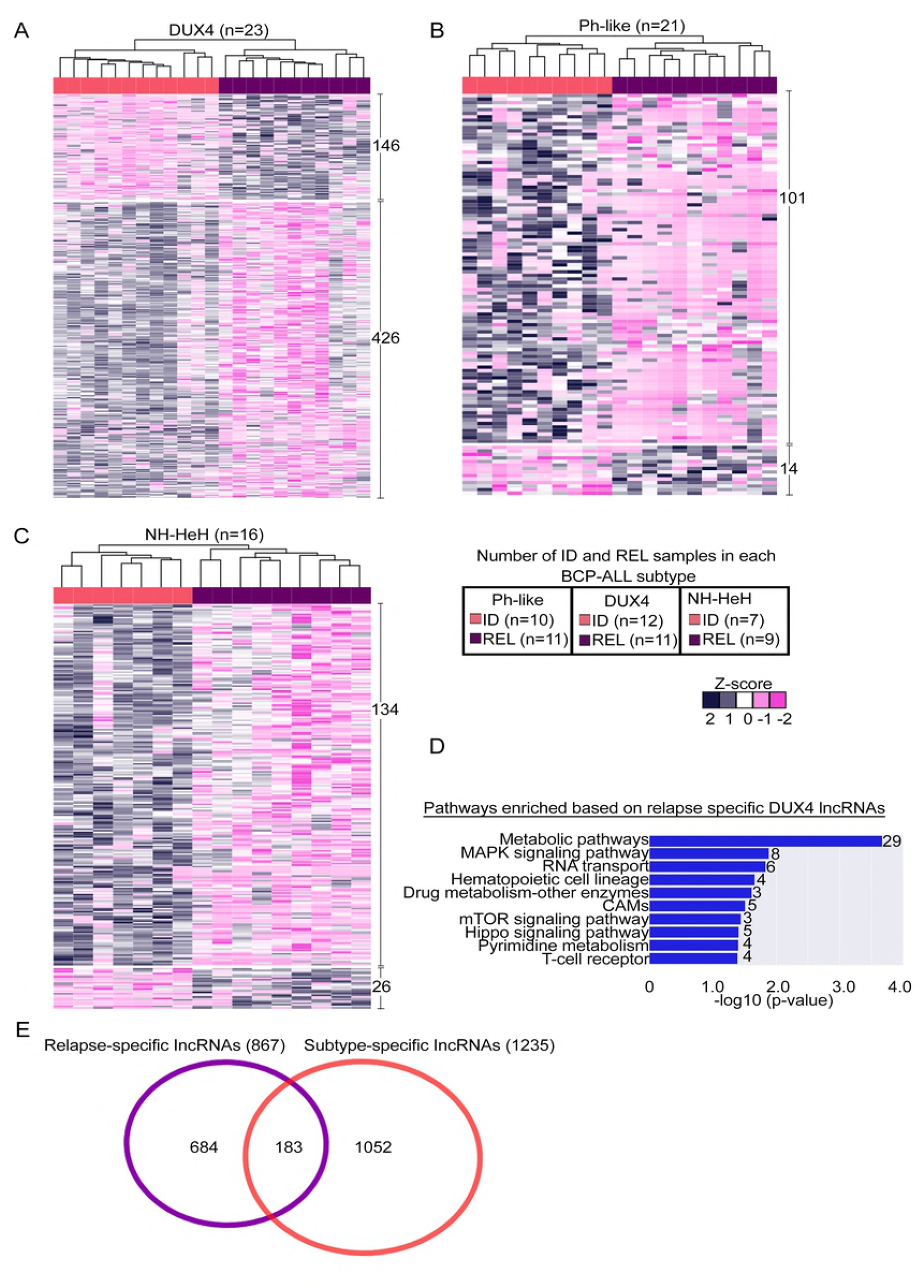
Relapse-specific DE lncRNAs from BCP-ALL subtypes. (A-C)Heatmap depicting the hierarchical clustering on relapse-specific DE lncRNAs signature on Z-score transformed LIMMA normalized expression values from DUX4, Ph-like and NH-HeH subtypes. Each heatmap shows the up and down regulated lncRNAs specific to ID and REL samples. (D) Molecular pathway analysis with the number of genes involved in each pathway from the enrichment analysis of the nearby (< 100 kb proximity) cis proteincoding genes correlated (Pearson correlation > 0.55 and P-value <= 0.05) with relapsespecific DE lncRNAs in the DUX4 subtype. The legend box indicates the number of ID and REL samples within each group. CAMs : Cell adhesion molecules. (E) The overlap between relapsespecific and subtype-specific lncRNAs from three subtypes.

### Subtype specific BCP-ALL lncRNAs show epigenetic alterations

For the analysis of the methylation status of loci located at the lncRNAs genomic position in the BCP-ALL subtypes, we used DNA methylation array data (collected from Illumina 450k methylation array) from the same patients (n = 46) including matched diagnosis (ID) and relapse (REL) samples (n = 82). The distribution of DNA methylation levels of CpG sites (n = 60,021, Table S4) associated with 7,160 lncRNAs was compared with CpG sites associated with PC genes across all BCP-ALL samples. Unlike the expression levels, the distribution of DNA methylation (hypo-methylation or hyper-methylation) between lncRNAs and PC genes were similar (Fig S1B). Given the robust separation of BCP-ALL subtypes on DNA methylation profile of CpGs associated with lncRNAs on the PCA analysis (Fig 4A), we next studied the differential hypo-methylated (methylation difference value < 0; *P*-value <= 0.05) and hyper-methylated (methylation difference value > 0.2; *P*-value <= 0.05) CpGs associated with lncRNAs in each subtype (see materials and method). The hierarchical clustering of differentially methylated (DM) lncRNAs showed distinct methylation patterns of each subtype, concordant with the DE lncRNAs signature (Fig 4B-D, Table S4). In the DUX4 and NH-HeH subtypes the number of hypo-methylated lncRNAs (differential methylation value < 0, *P*-value <= 0.05) were higher compared to the number of hyper-methylated lncRNAs. We classified the DM lncRNAs based on their genomic regions as gene body methylated and promoter-TSS methylated. In the promoter methylated lncRNAs we identified the same trend with high degree of hypo-methylated and lower number hyper-methylated lncRNAs in DUX4 and NH-HeH subtypes. However, the Ph-like subtype has shown a higher degree of hyper-methylated DM lncRNAs than hypo-methylated DM lncRNAs. The list of subtype-specific DM lncRNAs from three subtypes contained previously defined epigenetically altered lncRNAs from other cancer types, for example, we observed the oncogenic lncRNAs *LINC00312* (40), *PVT1*, and *TCL6* (41), which are differentially methylated in at least one of the three subtypes. Together, this data illustrates that epigenetically altered lncRNAs within three BCP-ALL subtypes.

**Fig 4:**
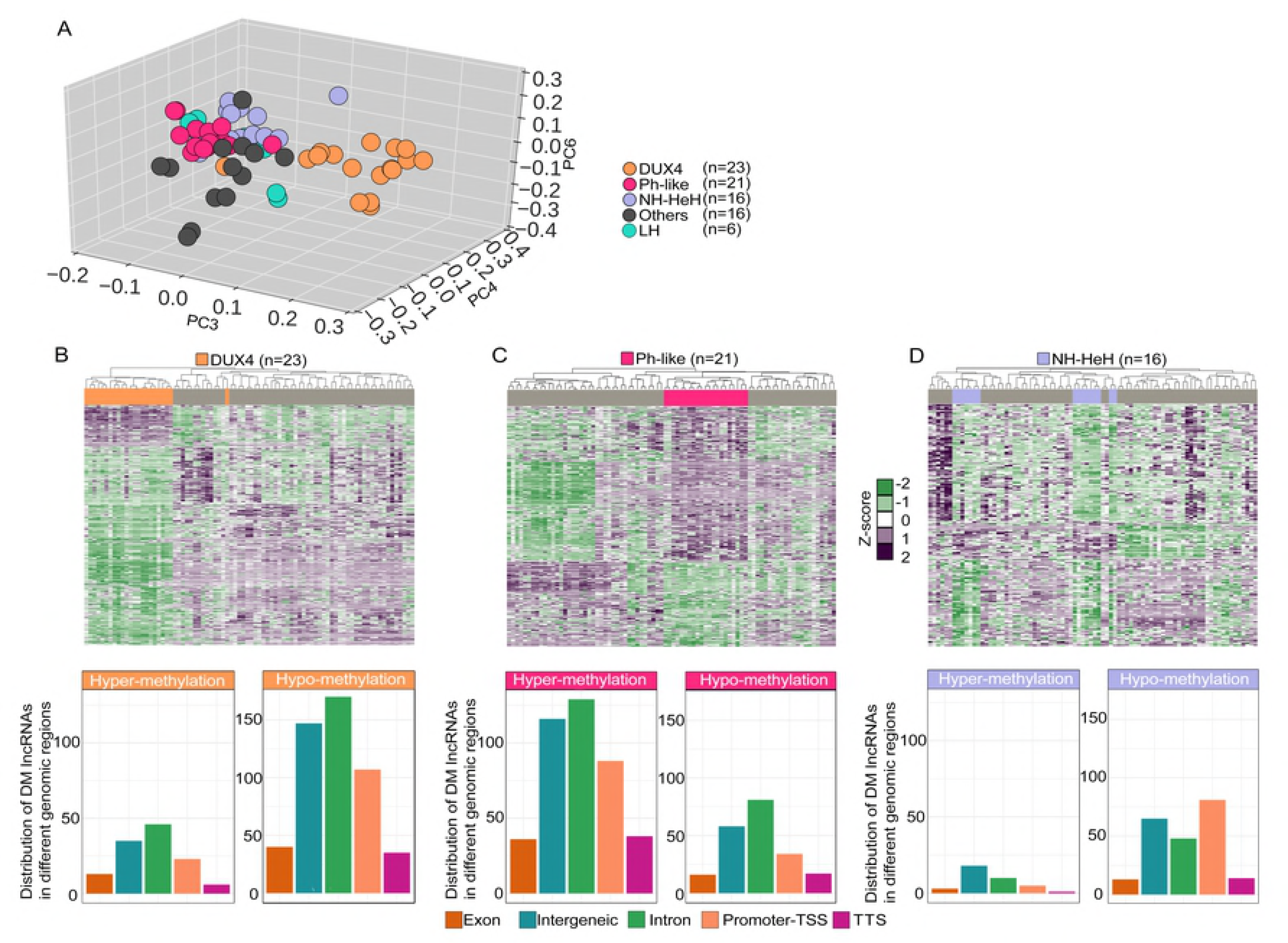
Hierarchical clustering of CGID’s associated with DM lncRNAs. (A) PCA of CpG’s associated with lncRNAs on SWAN normalized β values on 82 BCP-ALL samples obtained from DNA methylation array. Each point represents a BCP-ALL sample. DUX4, Ph-like, NH-HeH, LH and others are represented by orange, rose, blue, green and gray, respectively. (B) The heatmap representing hierarchal clustering on 544 differentially methylated (DM) CGID’s associated with 434 lncRNAs in DUX4 subtype. In the DUX4 subtype, we identified 328 (76%) differentially hypo-methylated and 106 (25%) hyper-methylated lncRNAs. (C) The heatmap representing hierarchal clustering on 518 DM CGID’s associated with 450 lncRNAs in the Ph-like subtype. In Ph-like subtype, we observed 302 (67%) hyper-methylated lncRNAs and 148 (33%) hypo-methylated lncRNAs. (D) The heatmap representing hierarchal clustering on 295 DM CGID’s associated with 234 lncRNAs in NH-HeH subtype. In the NH-HeH subtype, we identified 200 (86%) hypo-methylated and 34 (14%) hyper-methylated lncRNAs. The heatmap is ploted using SWAN normalized beta values. The barplots below each heatmap represents the distribution of DM lncRNAs in the genome (Promoter-TSS and gene body) lncRNAs from each subtype. The distribution DM Promoter-TSS lncRNAs are as follows: 25%, 29% and 39% in DUX4, Ph-like, and NH-HeH subtype, respectively.

### Correlation between differentially expressed and differentially methylated lncRNAs

In order to define whether the subtype-specific promoter methylation impacts on the expression level, we compared the promoter-TSS differential CpG methylated lncRNAs (n = 227) with its differential expression signature. We observed 44 lncRNAs with differential methylation pattern in their promoter region with differential expression pattern at RNA level. Out these, lncRNAs harboring significant hypo-methylation and hyper-methylation pattern (Pearson correlation, 2-tailed *P*-value <= 0.05) at the promoter region accounted for 23 (Table 4) lncRNAs across the three BCP-ALL subtypes.

**Table 4:**
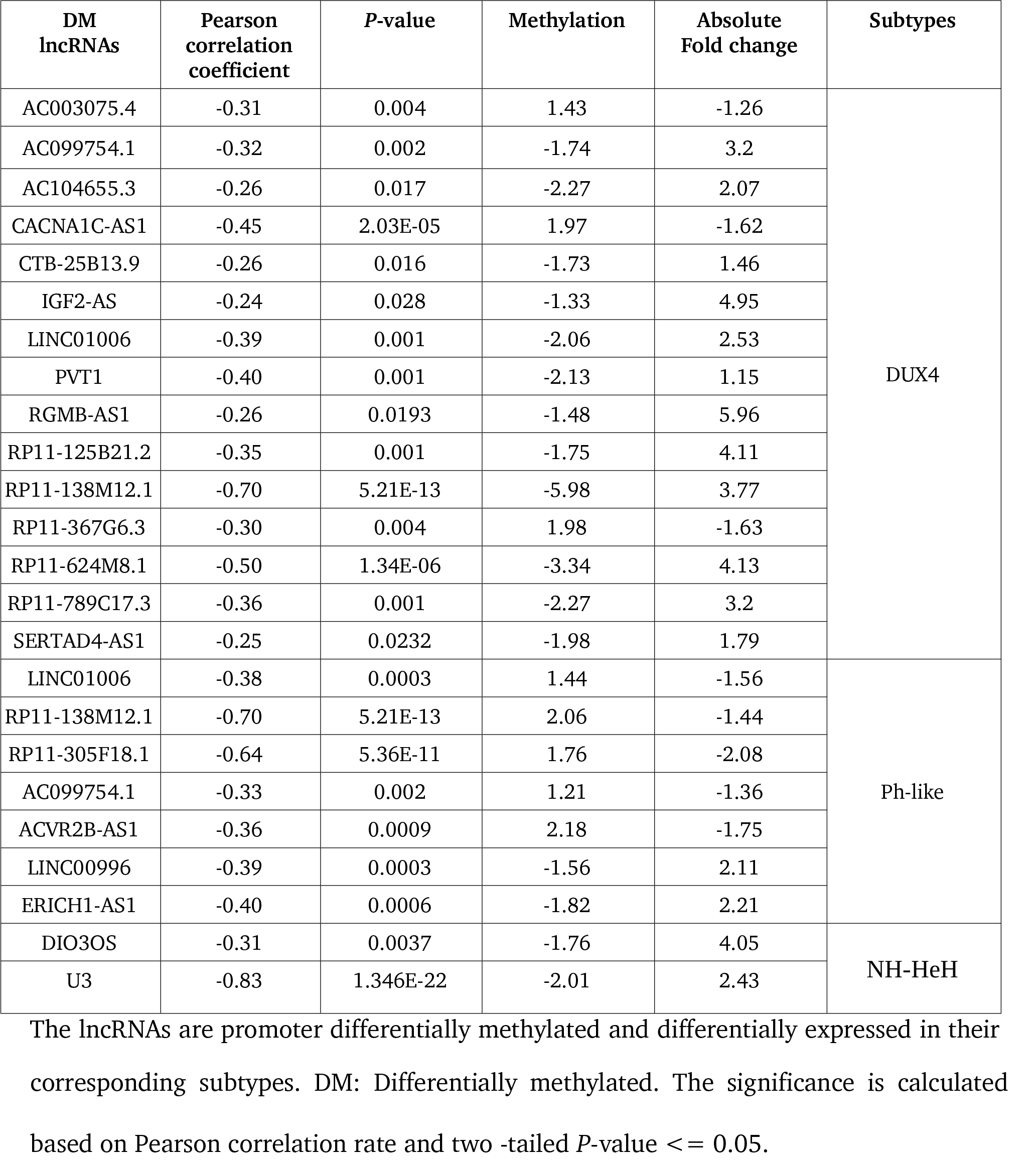
The list of significantly correlated DNA methylation and expression for promoter methylated lncRNAs (n = 23) from BCP-ALL subtypes.

Of these 23 putative epigenetically facilitated lncRNAs, 15 were related to the DUX4 subgroup (Fig 5A) including the novel lncRNAs, *R11-138M12.1* and *RP11-624M8.1*. These were significantly hypo-methylated at their promoter region and transcriptionally up-regulated in the DUX4 subgroup (Pearson correlation coefficient = −0.69; *P*-value = 5.1E-13 for *R11-138M12.1*; Pearson correlation coefficient = −0.50; *P*-value = 1.36E-06 for *RP11-624M8.1*; Fig 5B and 5C). In the Ph-like subtype, we observed 7 lncRNAs with promoter methylation (Fig 5D); interestingly, the same lncRNA *R11-138M12.1* showed significant hypermethylation at the promoter region and a concordant down-regulation in the Ph-like subgroup (Fig 5E). Besides these novel lncRNAs, we identified lncRNAs previously reported in the context of different cancers from our epigenetically altered results. The lncRNAs *PVT1* (Pearson correlation coefficient = −0.40, 2-tailed *P*-value <= 0.001), and *DIO3OS* (42) (Pearson correlation coefficient = −0.31, 2-tailed *P*-value = 0.0037) are examples, which we observed in the DUX4 and NH-HeH subtype with significant anti-correlation (2-tailed *P*-value <= 0.01) to its expression level. Around 46% (n = 512) of DM subtype-specific lncRNAs are localized in the intronic and intergenic genomic regions. We next aimed to investigate whether these lncRNAs regions has chromatin markers encoded within their genomic location. Recent human genome-wide chromatin marker study (43) has provided us with a rich resource to identify chromatin markers. Genome-wide mapping of B-lymphocyte cell line by searching for epigenetic markers within our DM subtype-specific intronic and intergenic regions revealed a significant number of lncRNAs (n=53) (Table S4, Fisher extract test P-value = 2.2E-16) with enchacer and insulator markers (Table S4). Out of these, lncRNAs, *RP11-134O21.1*, *RP11-398B16.2*, *RP11-689B22.2*, *CTC-458I2.2* and *LINC00880* were DE expressed, with a significant negative correlation between DNA methylation and expression levels in the DUX4 subtype (Table 5).

**Fig 5:**
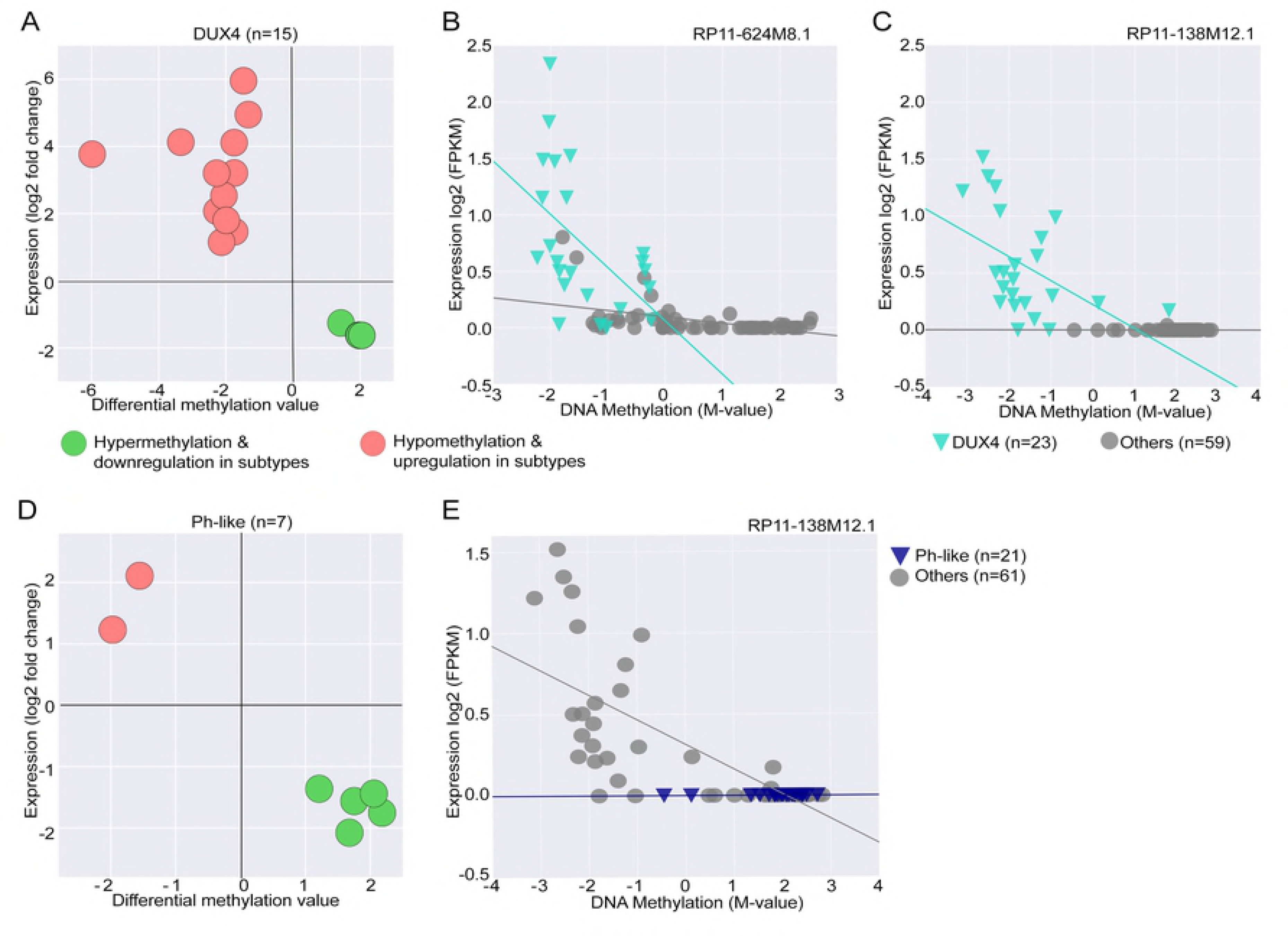
The epigenetically altered promoter methylated lncRNAs and their expression. (A) The promoter methylated lncRNAs with significant negative correlation with DE expression profile from the DUX4 subtypes. (B-C) Two representative examples of hypo-methylated lncRNAs with increased expression profile from DUX4 subtype. LncRNAs, *RP11-138M12.1* (Pearson correlation coefficient = −0.69, 2-tailed *P*-value = 5.21e-13), *RP11-624MB.1* (Pearson correlation coefficient = −0.50, *P*-value = 1.36e-06) are examples with hypo-methylation and up-regulated expression pattern with significant inverse correlation between DNA methylation and expression levels. (D) The promoter methylated lncRNAs with significant negative correlation with DE expression profile from the Ph-like subtypes. (E) A representative example of the promoter hyper-methylated lncRNA, *RP11-138M12.1* (Pearson correlation coefficient = −0.69, 2-tailed *P*-value = 5.21e-13) with down-regulated expression pattern, and with inverse correlation within the Ph-like subtype.

**Table 5:** The list of significantly correlated DNA methylation and expression for intronic and Intergenic methylated lncRNAs (n = 5) from DUX4 BCP-ALL subtypes.

| DM lncRNAs | Absolute Fold change | Methylation value | Pearson correlation rate | P-value | Epi-markers | Biotype |
| --- | --- | --- | --- | --- | --- | --- |
| RP11-134O21.1 | 2.54 | -1.56 | -0.63 | 1.9E-010 | Enhancer | Intron |
| RP11-398B16.2 | 2.08 | -1.85 | -0.47 | 0.0007 | Insulator |  |
| RP11-689B22.2 | 1.52 | -3.37 | -0.47 | 0.008 | Enhancer |  |
| CTC-458I2.2 | -1.16 | 3.38 | -0.42 | 0.0001 | Enhancer |  |
| LINC00880 | -1.45 | 2.23 | -0.25 | 0.02 | Enhancer | Intergenic |

The significance is calculated based on Pearson correlation rate and two -tailed *P*-value $\leq 0.05$ . The lncRNAs are promoter differentially methylated and differentially expressed in their corresponding subtypes. These lncRNAs are with enhancer and insulator epigenetic markers. DM: Differentially methylated.

These findings suggest that epigenetic silencing and activation of promoter lncRNAs may be a mechanism that contributes to the dysregulation of expression of lncRNAs. In addition to that, both intronic and intergenic DM lncRNAs associated with strong enhancer and insulator regions can accelerate its expression at the epigenetic level.

### Epigenetic alterations of subtype-specific lncRNAs are associated with evelvated expression of tumor genes located at their *cis* region

We next investigated the relationship between the epigenetic alterations of DM subtype-specific lncRNAs (n = 1118) and the aberrant expressions of their *cis* PC genes. We found 78 protein-coding genes located in their *cis* region, out of these 33 protein-coding genes have shown a significant up-regulated and down-regulated expression pattern in their corresponding subtypes (Table 6).

**Table 6:** The list of DNA methylated lncRNAs and differentially up and down-regulated *cis* PC genes from three subtypes.

| Subtype-specific Methylated lncRNAs | DNA methylation | Cis PC | Absolute Fold change | Subtype |
| --- | --- | --- | --- | --- |
| CTD-2231H16.1 | -1.6803385981 | **PLEKHG4B** | 2.4840131576 | DUX4 |
| RP11-80H8.4 | -4.8712973818 | **CHST2** | 5.021248353 |  |
| IGF2-AS | -1.5149692084 | **IGF2** | 6.7074586227 |  |
| RP11-332H18.4 | 1.7840373497 | **TBX2** | 5.6459057649 |  |
| RP11-332H18.4 | 1.7840373497 | TBX4 | 1.5056957579 |  |
| RP11-624M8.1 | -3.3400278742 | HEY2 | 3.5653881965 |  |
| CTB-25B13.9 | -1.7341024339 | REEP6 | 1.0135168425 |  |
| RGMB-AS1 | -1.4711332705 | **RGMB** | 4.4287098777 |  |
| AC073316.1 | -2.1346425435 | **SDK1** | 4.2539854375 |  |
| CTB-35F21.2 | -3.3996007483 | CXXC5 | 1.5322350503 |  |
| CTB-35F21.2 | -3.5262894443 | PSD2 | 2.0348265047 |  |
| AC099754.1 | -1.7432607805 | LRRC3B | 3.2956570803 |  |
| AC078883.4 | -3.4089445499 | CORO1C | 0.797282079 |  |
| AC078883.4 | -3.4089445499 | **ITGA6** | 3.2489394471 |  |
| RP11-314O13.1 | -1.5432811428 | CDYL2 | 1.7070802978 |  |
| RP3-455J7.4 | -3.0695619042 | CREG1 | 1.5618184051 |  |
| RP11-125B21.2 | -1.749750681 | VLDLR | 4.2858349208 |  |
| RP11-676J12.8 | -1.8927665898 | GLOD4 | 0.6471353534 |  |
| LINC00114 | 3.4320004844 | ETS2 | -0.7176247124 |  |
| RP11-367G6.3 | 1.9798401851 | THBS4 | -1.5796823436 |  |
| CTC-458I2.2 | 3.3822300323 | CTGF | -1.892559675 |  |
| RP11-69I8.3 | 2.355315514 | AHR | -2.4815111685 |  |
| RP11-293A21.1 | -3.6963992484 | STIM2 | 2.487452138 | NH-HeH |
| LINC00114 | -2.9731205546 | ETS2 | 0.9908037963 |  |
| ACVR2B-AS1 | 2.3094716973 | ACVR2B | -2.4697071313 | Ph-like |
| AGAP1-IT1 | 2.0926102582 | AGAP1 | -2.9857451706 |  |
| RP11-69I8.3 | -3.1689478827 | CTGF | 1.8492946882 |  |
| LINC00996 | -1.553999019 | GIMAP8 | 1.4145369937 |  |
| AC099754.1 | 1.2121139088 | LRRC3B | -2.0406367867 |  |
| CTB-79E8.2 | -2.2285747372 | NEURL1B | 1.1469246971 |  |
| AL133493.2 | 2.0838061516 | PCBP3 | -3.0795988736 |  |
| RP11-420G6.4 | -2.4300037457 | SERPINB1 | 0.5848645847 |  |

The table represents hypo-methylated and hyper methylated lncRNAs from three subtypes elevating and diminishing the expression of the protein-coding genes localized at their cis regions. The PC genes differentially up and down regulated in the respective subtypes include tumor genes as well. The highlighted rows are some examples of oncogenes with up-regulated expression profile within DUX4 subtype.

Intriguingly, the up-regulated PC genes include known tumor genes from various cancer types. For example, *IGF2* (44) (absolute fold change = 6.70, *adj.P*-value = 0.0061), *CTGF* (45) (absolute fold change = 1.85, *adj.P*-value = 0.02) and *ETS2* (46) (absolute fold change = 0.99, *adj.P*-value = 0.01) from DUX4, Ph-like and NH-HeH subtypes respectively (Fig 6 A-C). Together, this illustrates a set of lncRNAs which are capable of epigenetically elevating and silencing the expression profile of tumor genes localized in its *cis* region in BCP-ALL subtypes.

**Fig 6:**
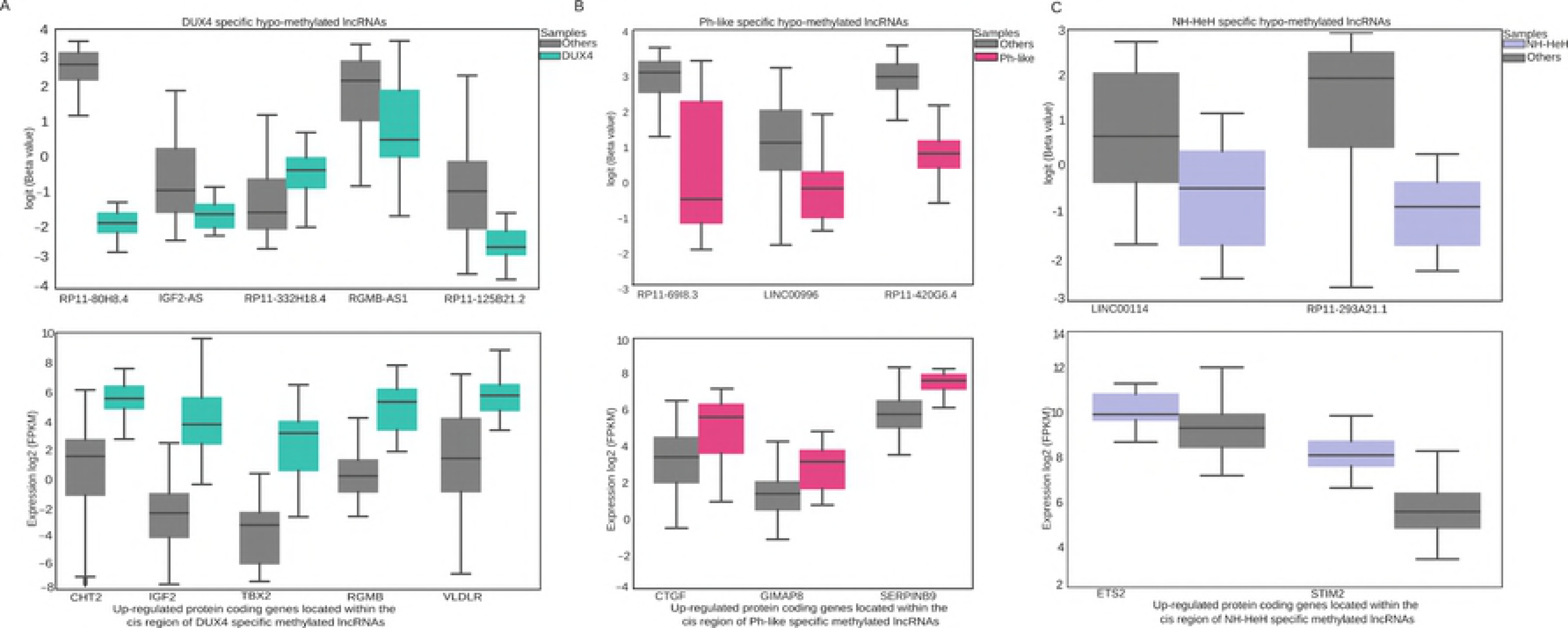
Differentially methylated lncRNAs epigenetically altered expression levels of the *cis* oncogenes. (A) The upper panel of boxplot represents the DNA methylated lncRNAs, the boxplot below that represents their corresponding *cis* ocogenes which are up-regulated in DUX4 subytpe. The barplox shows representative examples of hypo-methylated lncRNAs, *RP11-80H8.4* (DNA methylation value = −4.87, *P*-value = 0.0001), *IGF-AS2* (DNA methylation value = −1.52, *P*-value = 0.011), *RP11-332H18.4* (DNA methylation value = 1.79, *P*-value = 0.0057), *RGMB-AS1* (DNA methylation value = −1.47, *P*-value = 0.007), and *RP11-125B21.2* (DNA methylation value = −1.75, *P-*value = 0.007) and its corresponding significatly up-regulated *cis* oncogenes, *CHT2* (absolute log fold change = 5.021, FDR = 2.39E-08), IGF2 (absolute log fold change = 6.71, FDR = 41.412E-15), *TBX* (absolute log fold change =5.64, FDR = 3.97E-13), and RGMB (absolute log fold change = 4.42, FDR = 3.02E-16) within DUX4 subytpe. (B) The upper panel of boxplot represents the DNA methylated lncRNAs, the boxplot below that represents their corresponding *cis* ocogenes which are up-regulated in Ph-like subytpe. The barplox shows representative examples of hypo-methylated lncRNAs, *RP11-69l8.3* (DNA methylation value= −3.16, P-value = 4.46E-08), *LINC00996* (DNA methylation value = −1.55, *P-*value = 0.02), *CTB-79E8.2* (DNA methylation value = −2.22, *P*-value = 0.009), *RP11-420G6.4* (DNA methylation value = −2.43, *P*-value = 0.004) and its corresponding significatly up-regulated *cis* oncogenes, *CTGF* (absolute log fold change = 1.85, = 0.02), *GIMAP8* (absolute log fold change = 1.14, FDR = 0.004), *NEURLB* (absolute log fold change = 1.14, FDR = 0.09), *SERPINB1* (absolute log fold change = 1.63, FDR = 0.0004) within Ph-like subtype. (C) The upper panel of boxplot represents the DNA methylated lncRNAs, the boxplot below that represents their *cis* ocogenes which are up-regulated in NH-HeH subytpe. The barplot shows representative examples of hypo-methylated lncRNAs, *LINC00114* (DNA methylation value = −2.97, P-value = 0.0003), *RP11-293A21.1* (DNA methylation value = −3.69, P-value = 0.00071) and its corresponding significatly up-regulated *cis* oncogenes, *ETS2* (absolute Fold change = 0.99, FDR = 0.019) and *STIM2* (absolute Fold change = 2.48, FDR = 6.42E-10) within NH-HeH subtype. False discovery rate: FDR

## DISCUSSION

Although previous studies have demonstrated the involvement of lncRNAs in acute leukemias (25,26) comprehensive characterization of the transcriptome, epigenetic regulation and functional contribution of lncRNAs in distinct BCP-ALL subtypes are lacking. lncRNAs, as the novel class of functional molecules involved in cancer biology, is defined in the stratification of different molecular subtypes in various cancers (47–49). However, their role in BCP-ALL subtypes has not been investigated. Here, we unravel the lncRNAs landscape using transcriptome and methylome data from 46 (adult and pediatric) relapsed BCP-ALL patients focusing on the three molecular subtypes namely, DUX4, Ph-like, and NH-HeH. Our integrated transcriptomic analyses using RNA-seq and DNA methylation brings significant insights and advances over other studies: it provides the most comprehensive novel datasets so far for BCP-ALL subtypes, a resource of subtype-specific and relapse-specific lncRNAs, potential lncRNAs functions and uncovers their epigenetic alterations of the BCP-ALL subtypes. We identified 1235 DE subtype-specific lncRNAs dysregulated in at least one of the three subtypes. Compared to the pan-cancer comprehensive set of aberrantly expressed lncRNAs we found 66% (712 out of 1564) of our DE lncRNAs were more specific for our subtypes introducing novel insight in the non-protein-coding part of the genome in BCP-ALL subtypes.

Another important aspect of our study is the identification of relapse-specific dysregulated lncRNAs across three BCP-ALL subtypes. A closer look into the relapse-specific lncRNAs signature identified lncRNAs previously described as oncogenic: lncRNAs including, *RP11-701P16.5 (50)*, *SLC38A3 (51)*, and *LINC00312 (40)*, which are up-regulated in relapsed samples within DUX4 subtype (Table 7).

**Table 7:** Previously reported lncRNAs identified as relapse-specific lncRNAs in BCP-ALL subtypes.

| Relapse-specific lncRNAs | Disease association |
| --- | --- |
| TCL6 (DUX4) | Chromosomal translocations T-cell leukaemia/lymphoma (2) |
| LINC00312 (DUX4, Ph-like, NH-HeH) | Proliferation, invasion, and migration of thyroid cancer, Nasopharyngeal carcinoma (3) |
| miR-17-92a-1 (DUX4, Ph-like, NH-HeH) | Development, progression, and aggressiveness of colorectal cancer (4) |

The differentially expressed lncRNAs between relapse (REL) and initial diagnosis (ID), from three subtypes, which were previously reported for its disease association, selected representative examples from relapse-specific lncRNAs, which are previously identified in other diseases.

Importantly, apoptosis suppressor lncRNA in myc-driven lymphomas (52) *miR-17/92* cluster host gene (*MIR17HG*) is up-regulated in relapse samples within the Ph-like subtype and down regulated in relapsed samples within DUX4 and NH-HeH subtypes. Overall, the relapse-specific lncRNAs highlights the oncogenic relevance in BCP-ALL subtypes.

Besides the oncogenic properties, lncRNAs can act as prognostic markers (53) and aid for disease diagnosis and treatment. A subset of our relapse-specific lncRNAs (n = 61, Table S3) is recently identified as prognostic markers in 14 Pan-Cancer data (36) types, including Lung Cancer Associated Transcript 1 (*LUCAT1*), which is previously reported for its drug resistance in solid cancer (54). Within the DUX4 subtype, we identified up-regulated expression of *LUCAT1* at relapse, potentially providing a novel insight into treatment resistance for BCP-ALL subtypes. Together, this illustrates the catalog of relevant lncRNAs in different subtypes of BCP-ALL as subtype-specific and relapse-specific markers with the potential of RNA based treatments in the treatment of BCP-ALL subtypes.

The dissection of the regulatory pathways mediated by the molecular subtype-specific and relapse-specific lncRNAs revealed the activation of pivotal signalling pathways across three BCP-ALL subtypes. The functional analysis using guilt-by-association approach highlights the subtype-specific and relapse-specific lncRNAs associated with activation of signaling pathways and metabolic pathways that are associated with leukemogenesis including TGF-Beta, hippo, P53, and JAK-STAT, cytokine-cytokine receptor, endocytosis, mTOR and metabolic pathways. Characterization of the lncRNAs involved in this pathway may potentially reveal novel targets in molecular therapies.

The functional insights of relapse-specific and subtype-specific lncRNAs revealed biological relevance to BCP-ALL subtypes including cell cycle functions, signal transduction, cell migration and metabolic processes. Some of the functions predicted here corroborate previous studies emphasizing the strengths of the employed guilt-by-association. For example, lncRNA *AC002454.1*, which we associated to the PIK3-AKT pathway in Ph-like subtype, was recently reported to regulate cyclin-dependent kinase (*CDK6*) to participate in cell cycle disorder (55). The *CDK6* gene appears to be frequently dysregulated in hematopoietic malignancies (39) and is hence attributed a critical role in tumorigenesis, also shown in ALL driven by mixed lineage leukemia fusion proteins (56). In Ph-like subtype, both *CDK6* and *AC002454.1* are correlated and up-regulated specifically in Ph-like samples, suggesting they displayed enhancer-like functions. We identified 8 relapse-specific lncRNAs (Table S3) associated with metabolic pathways in the DUX4 subtype overlapping with lncRNAs reported (57) to synergistically dysregulate metabolic pathways in multiple tumour context.

Besides known lncRNAs, we also identified novel lncRNAs associated with activation of key signalling pathways. For instance, in DUX4 subtype, we identified a set of novel lncRNAs associated with TGF-beta pathway, including the antisense *RP11-224019.2*, with a significant positive correlation to the *TGFB* gene. Recently, there are a number of lncRNAs documented to be associated with TGFß signalling pathway, including MEG3 regulating the TGFB2 pathway in breast cancer (58). However, lncRNAs associated with the TGFß pathway in BCP-ALL subtypes have not been reported. The co-expression of *RP11-224019.2* and *TGFB* in DUX4 subtype may indicate their functional relatedness or regulatory relationships. In addition to that, a notable observation was a strong correlation between relapse-specific lncRNAs with genes involved in the activation of metabolic pathways in the DUX4 subtype. We identified 112 relapse-specific lncRNAs co-expressed with 29 (Table S3) PC genes activated in metabolic pathways, including previously reported 8 biomarker lncRNAs. For example, we identified oncogenic lncRNA *LUCAT1* reported to be associated with poor prognosis in lung cancer (54). However, the *LUCAT1* has not yet been reported in the BCP-ALL context. The global co-expression analysis and gene-expression profiling suggest important and previously unappreciated roles of lncRNAs in the BCP-ALL subtypes. Our analyses provide important functions of subtype-specific and relapse-specific lncRNA genes whose expression correlates tightly with oncogenic coding genes.

Although we observed that subtype-specific lncRNAs and subtype-specific protein-coding genes were predicted to activate or inhibit the same pathways, some important exclusivity was observed. For instance, the signalling pathways such as the PI3K and mTOR in Ph-like subtype was enriched only in the lncRNAs based enrichment analysis, whereas these pathways did not appear to be enriched/dysregulated in the mRNA based analysis. The PI3K and mTOR signalling pathways control proliferation, differentiation, and survival of hematopoietic cells (59). Consistent with our studies, other studies indicated the potency of lncRNAs facilitating the cancer cell growth through mTOR and PI3K signalling pathways (36,47,60) yet reports on BCP-ALL subtypes are lacking. Considering the functional nexus between Ph-like specific lncRNAs and the activation of pathways such as mTOR and PI3K signalling pathways, targeting those lncRNAs may be a promising novel therapeutic target for BCP-ALL subtypes.

Our work additionally underscores the importance of epigenetic alterations in modulating lncRNAs transcriptional activities. Although previous studies (16, 57) have demonstrated cross-talk between DNA methylation and transcriptional activities of lncRNAs, their role in the etiology of BCP-ALL subtypes has not been investigated. DNA methylation analyses of lncRNAs revealed that DNA methylation might underlie the differential expression of BCP-ALL subtype-specific lncRNAs. Subtype-specific lncRNAs identified here have been reported by previous studies. For example, *SOX2-OT* (62), *LINC00312* (63), *TCL6* and *PVT1*, are onco-lncRNAs, which are promoter methylated in one of the three subtypes. The lncRNA, *PVT1* was reported for its MYC activity (64,65) and as oncogenic lncRNA with multiple roles in cell growth, dysfunction, and differentiation in AML (66). Both lncRNAs, *LINC00312* and *TCL6* have been extensively investigated on expression levels but not on the epigenetic level. The promoters of both *TCL6* and *LINC00312* were observed to be hyper-methylated with corresponding diminished expression in the DUX4 and NH-HeH samples. Notably, the DNA methylation analysis of lncRNAs revealed that DNA methylation might underlie the differential expression of subtype-specific lncRNAs. Our analysis identified 23 subtype-specific lncRNAs showing hypo-methylation and hyper-methylation pattern at their promoter region that are significantly correlated with their diminished and increased expression in respective subtypes. In addition to that, we have identified 33 epigenetically co-regulated oncogenes localized in the *cis* regions of hypo-methylated and hyper-methylated lncRNAs from three subtypes. Interestingly, the oncogene associated with leukemia, IGF2 (44) has shown an elevated expression level in Ph-like subtypes corresponding to the hypo-methylation of its antisense, IGF2-AS1. These findings suggest that epigenetic silencing of lncRNAs genes may be a mechanism that contributes to the dysregulation of expression of lncRNAs and their *cis* genes in BCP-ALL subtypes.

Overall, our study provides an in-depth analysis of the lncRNA transcriptome and epigenome in BCP-ALL subtypes and provides numerous new lncRNAs markers associated with subtype and relapse-specificity and with epigenetic alterations in BCP-ALL subtypes. Additionally, we also demonstrated these lncRNAs might contribute to the regulation of key signalling pathways involved in BCP-ALL. In summary, our study provides a comprehensive set of dysregulated lncRNAs from BCP-ALL subtypes derived using different integrative approaches. This can serve as a major resource of BCP-ALL subtype-specific lncRNAs and their mechanisms of action in detail that might pave the way for the future studies to investigate key biomarkers and potential therapeutic targets in BCP-ALL subtypes.

## Materials and Methods

### Patient samples

Patients (n = 46) used in this project were negatively selected for fusion genes detectable by routine diagnostic workup (BCR-ABL, MLL translocations, ETV6-RUNX1) from 26 pediatric and 22 adult patients. From these patients we collected 44 samples at initial diagnosis (ID) and 44 samples with relapse (REL). All patients were treated in population-based German study trials (GMALL for adult and BFM for pediatric patients). A written informed consent to participate in these trails according to the Declaration of Helsinki was obtained from all patients. The studies were approved by the ethics board of Charité, Berlin.

### Overview of RNA-seq and DNA methylation array data

To generate transcriptome profiles of patient samples, mRNA was isolated by using Trizol reagent (Life Technologies, Grand Island, NY) procedure from the bone marrow mononuclear cells (MNCs) of the ID and REL samples. The paired-end RNA sequencing was done on Illumina HiSeq4000 platform (multiplexing) in the high throughput sequencing core facility, German Cancer Research Center, Heidelberg, Germany. The RNA-Seq was performed by using six samples per lane, which resulted in an average of 64 Million mapped paired reads per sample. For methylation, genomic DNA was isolated using unstranded Allprep extraction (Qiagen, Hilden, Germany) from the bone marrow of same patients (ID and REL samples) was then hybridized onto an Illumina Infinium HumanMethylation450K. From the DNA methylation chip we identified 60,021 probes annotated to 7,190 lncRNAs.

### RNA-seq read alignment and quantitative extraction

The STAR aligner (version 2.4.0.1) (67) (2-pass alignment parameters) was used to align paired-end reads to the human genome reference. The human genome reference files used for processing RNA-seq samples were the hg19 (GRCh37) genome version for alignment and transcript annotation from GENCODE version 19 (equivalent Ensembl GRCh37). The transcriptome construction and gene-level counts for each sample were obtained using StringTie (68). The read count information from the files generated by StringTie was extracted using the “prepDE.py” python script provided by the StringTie. We detected 84% of 13,860 lncRNAs (including 23,898 transcripts) annotated by GENCODE (V19) from our samples (FPKM > 0 for multi-exon lncRNAs and FPKM > 0 for single exonic lncRNAs) showing that our sequencing depth was good.

### Sample clustering and differential expression analysis for subtype-specific and relapse-specific lncRNAs

We performed PCA using the *prcomp* R function on 13,860 lncRNAs from RNA-seq and 60,021 CpG’s on lncRNAs from DNA methylation datasets. The PCA plots were plotted using python matplotlib axes3D function. The R bioconductor package Linear Models for Microarray (*LIMMA) Voom* (69) was used on gene-level expression data for identifying the subtype-specific and relapse-specific differentially expressed (DE) lncRNAs. The subtype-specific DE lncRNAs were identified by implementing separate design matrix for the three subtypes (DUX4, Ph-like and NH-HeH). Within each subtype, we used using all subtype samples versus the rest of the cohort. Within our cohort (82 samples from 46 patients), not all patients had matching ID and REL samples, and moreover, we wanted to compare across subtypes. LIMMA voom leveraged the sample imbalances and confounder (patient and samples) with its *duplicatecorrelation* function. We implemented *duplicatecorrelation* function which addressed all patient effects by estimating correlations of multiple samples from the same patient while allowing us to compare across the subtypes. Additionally, we included the ID and REL time factors into the design (*makeContrasts*) to avoid the inflation of the variance due to time factor for each subtype. The relapse-specific DE lncRNAs within each subtype were identified by analysing DE lncRNAs ID versus REL samples within each subtype separately. The significant DE genes were assigned based on the p-value < 0.01 and Fold change of >= +-1.5. The lncRNAs from GENCODE version 19 (equivalent Ensembl GRCh37) were used as reference annotation. The heatmaps and correlation based (Spearman method) hierarchical clustering of DE lncRNAs were performed on z-score transformed LIMMA normalized gene expression values using the R Bioconductor package ComplexHeatmap.

### Differential methylation data analysis

The ID and REL samples from the same patients have been assayed with the Illumina 450k methylation array. All the beta values have been normalized using the Subset-quantile Within Array Normalization (SWAN) method. In order to detect differentially methylated regions, we used the R package bumphunter (70) using the most variant quartile of the CpG probes. Bumphunter searches for differentially methylated regions in an annotation-unbiased manner. Separate bumphunter runs have been performed for ID and REL samples for every three subtypes (DUX4, Ph-like, and NH-HeH), using all subtype samples versus the rest of the cohort. We associated the differentially methylated regions from three BCP-ALL subtypes using HOMER (hypergeometric optimization of motif enrichment) suite of tool with the reference file GRCh37.74, using the -gene parameter. HOMER provided us with annotation of each probe, we separated lncRNAs from the output. The genomic regions were divided into promoter (+-2kb from transcription start site, TSS) and gene body. The gene body was defined if the CpGs were annotated in exonic, intronic or transcription termination site (TTS). The regions mapped to lncRNAs were then used for analysis. The significantly differentially hyper-methylated (Methylation difference value >= 0.2; *P*-value <= 0.05) and hypo-methylated (Methylation difference value =< 0; *P*-value <= 0.05) regions were used for further analysis. The intronic and intergenic differentially methylated (DM) lncRNAs were then mapped using ‘BedTools’ with the B-lymphocyte cell line “wgEncodeBroadHmmGm12878HMM.bed” in order to find the epigenetic markers. The significance of enrichment was calculated using Fisher’s exact test. The epigenetically altered lncRNAs were assigned if promoter methylated lncRNAs were differentially expressed and their DNA methylation values (log-transformed Beta values) and expression values (log-transformed FPKM values) are correlated. The most significant correlations (Pearson correlations coefficient, 2-tailed *P-*value <= 0.05) were classified later called as epigenetically altered lncRNAs.

### Functional predictions using guilt-by-association approach

In our study, we used the “guilt-by-association” (71) approach by establishing the pairwise expression correlations between DE lncRNAs (from all BCP-ALL subtypes) and it’s *cis* and *trans* protein-coding (PC) genes to predict the functions of subtype-specific lncRNAs. We located the *cis* and *trans* PC genes of DE lncRNAs using the GREAT tool (version v3.0.0) (72). All PC genes from GENCODE v19 annotation (n = 20698) were used in the analysis. The individual *cis* and *trans* genes for each DE lncRNAs were located within a genomic window of 100 kb and greater >100kb respectively. From each dataset, we then computed the pairwise expression correlation using Pearson correlation method between each lncRNAs and its *cis* and *trans* coding gene. The significantly co-expressed PC genes (Pearson correlation coefficient >= 0.55 and 2-tailed *P*-value <= 0.05) were further used for functional enrichment analysis using GeneSCF v1.0 (73). The functional enrichment analysis was performed using the KEGG database with a background of all protein-coding genes from GENCODE v19 (20,345). The functional terms were considered significant only if it is enriched with *P*-value <= 0.05.

## ACKNOWLEDGEMENT

We are grateful to Johanna Angermaier for discussions and Ulf Leser for the suggestions in data analysis.

## Author Contributions

The project was conceived and designed by: ARJ, CDB, MN, AA. ARJ developed bioinformatics pipeline and analyzed RNA-seq data. MN normalized the methylation data. MSP performed DNA methylation data analysis. MPS, LB, MN and CDB performed the analyses of clinical and molecular data. JOT, CS and KI performed the sample preparation. CMT, MAR, MB, NG, RKS, AVS, MS, MH, TB, SS, HS, SG, RKS, CE were involved in the sample collection, genetic characterization and provided molecular diagnostic data. All authors were involved in writing and reviewing the manuscript.

## Supporting Information caption

**Fig S1: The distribution of lncRNAs and PC gene expression and DNA methylation levels across samples.** (A) The level of distribution of expression between 13460 lncRNAs and 20,135 PC genes across 82 BCP-ALL samples. (B) The level of distribution of DNA methylation rate between 60,022 CpGs probes associated with lncRNAs region and 120,000 CpGs probes associated with PC genes across 82 BCP-ALL samples.

**Fig S2: BCP-ALL subtype-specific differentially expressed lncRNAs** (A-C) The hierarchical clustering representing lncRNAs clustering and expression differences of the compared subtypes DUX4, Ph-like and NH-HeH; corresponding to 736, 383, and 445 subtype-specific DE lncRNAs in DUX4, Ph-like and NH-HeH subtypes, respectively. In the DUX4 subtype, 100% of samples clustered together based on the DE lncRNAs signature. The hierarchical clustering of the subtype-specific DE lncRNAs revealed that 90% (19 out of 21 samples) of Ph-like samples clustered within the predefined Ph-like subtype. For the NH-HeH subtype 69% (11 out of 16 samples) of samples correlated and clustered together using the respective DE lncRNA signature. The BCP-ALL samples box representing the number of samples within each subtypes and versus (vs) the other samples used as control group in DE analysis. (D) The overlap between DE subtype specific lncRNAs from three subtypes versus public list of dysregulated lncRNAs from 12 different cancer types comprehensive cancer genome (CGC)

**Fig S3: Comparison of molecular pathways from cis and trans based analysis on subtype-specific DE lncRNAs.** A) Molecular pathway analysis from functional enrichment analysis on *trans* (>= 100 kb) protein-coding genes correlated (Pearson correlation coefficient >= 0.55 and two-tailed *P-*value <= 0.05) with DE lncRNAs in DUX4 subtype. (B) The molecular pathways overlapped between *cis* (< 100 kb proximity) and *trans* (> 100 kb) based functional enrichment analysis in the DUX4 subtype. (C) Molecular pathway analysis from functional enrichment analysis on *trans* (> 100 kb) protein-coding genes correlated (Pearson correlation coefficient >= 0.55 and two-tailed *P*-value <= 0.05) with DE lncRNAs in Ph-like subtype. CAMs : Cell adhesion molecules, CML: Chronic myeloid leukemia, AML: Acute myeloid leukemia.

**Fig S4: The subtype-specific lncRNAs co-expressed with oncogenes involved in key signaling pathways in DUX4 and Ph-like subtypes.** (A-B) Antisense *RP11-224O19.2* (absolute Fold change = 2.786, P-value = 9.74E-08) and its *cis* oncogene *TGFB2* (absolute Fold change = 3.84, P-value = 2.74E-10) are significantly up-regulated in DUX4 samples. (C) Antisense lncRNAs *R11-536K7*.5 located at *cis* region of oncogene *IL2RA*. Expression of antisense lncRNA *RP11-536K7.5* showed significant co-expression with expression of its *cis* oncogene *IL2RA*. Both *RP11-536K7.5* (absolute Fold change = 2.79, *P*-value = 3.07E-008) and *IL2RA* (absolute Fold change = 3.11, *P*-value = 3.97e-1) are up-regulated in Ph-like samples. (D) The expression of *cis* antisense lncRNA *AC002454.1* significant co-expressed with its *cis* oncogene *CDK6* in Ph-like subtype. Both *CDK6* (absolute Fold change = 1.01, *P*-value = 0.0005) and antisense lncRNA *AC002454.1* (absolute Fold change = 1.79, *P*-value = 0.00015) are up-regulated in Ph-like samples.

**Table S1: RNA-seq data used for analysis and subtype-specific lncRNAs from three subtypes**

**Table S2: The functionally involved subtype-specific lncRNAs from DUX4 and Ph-like subtypes. The trans and *cis*-acting subtype-specific lncRNAs**

**Table S3: The relapse-specific lncRNAs from three subtypes. The lncRNAs involved in functions from DUX4 subtype**

**Table S4: DNA methylation array dataset. The differentially methylated lncRNAs from three subtypes. List of cis-acting epigenetically active lncRNAs.**

## AVAILABILITY OF DATA AND ACCESSION NUMBERS

All sequencing data used in this study is available at the European Genome phenome Archive (accession number to be provided after acceptance of the manuscript)

## FUNDING

This work was supported by the German Cancer Aid (Deutsche Krebshilfe) [grant number 111533].

